# FSTL3 is a biomarker of poor prognosis and is associated with immunotherapy resistance in ovarian cancer

**DOI:** 10.1101/2024.12.10.627747

**Authors:** Maeva Chauvin, Estelle Tromelin, Julien Roche, Hyshem H Lancia, Marie-Charlotte Meinsohn, Caroline Coletti, Ngoc Minh Phuong Nguyen, Virginie Lafont, Henri-Alexandre Michaud, Ranjan Mishra, Laurent Gros, Nathalie Bonnefoy, David Pépin

**Author notes:** Corresponding authors: David Pépin, Pediatric Surgical Research Laboratories, Massachusetts General Hospital; Department of Surgery, Harvard Medical School, Boston, MA 02114. contributed equally Maeva Chauvin, Pediatric Surgical Research Laboratories, Massachusetts General Hospital; Department of Surgery, Harvard Medical School, Boston, MA 02114. The authors declare no potential conflicts of interest.

## Abstract

High-grade serous ovarian carcinoma (HGSOC), is associated with high mortality rates due to late-stage diagnosis and limited treatment options. We investigated the role of FSTL3 in ovarian cancer progression both as a prognostic biomarker and as a potential therapeutic target.

We measured levels of follistatin (FST) and follistatin-like 3 (FSTL3) in 96 ovarian cancer patient ascites samples and found that FSTL3 overexpression was more predominant than FST and associated with poorer survival outcomes. Mice implanted with an HGSOC syngeneic cell line bearing common alterations in ovarian cancer (KRAS^G12V^, P53^R172H^, CCNE1^oe^, AKT2^oe^) had increasing levels of FST and FSTL3 in serum during tumor growth. Further alteration of this model to generate a knockout of FST (KPCA.FSTKO) and an overexpression of human FSTL3 (KPCA.FSTKO_hFSTL3), revealed that FSTL3 expression was associated with a more fibrotic tumor microenvironment, correlating with an increased abundance of cancer-associated myofibroblasts (myCAFs), and cancer cells with a more mesenchymal phenotype. Tumors overexpressing FSTL3 had less immunocyte infiltration and a significantly reduced intratumoral T-cell abundance (CD4+, CD8+). FSTL3 overexpression completely abrogated tumor response to PPC treatment (Prexasertib combined with PD-1 and CTLA-4 blockade) compared to controls, suggesting that FSTL3 may be involved in immunotherapy resistance.

In conclusion, this study suggests a role for FSTL3 as a prognostic marker and as therapeutic target in HGSOC, where it may play a role in promoting a mesenchymal tumor phenotype, maintaining an immunosuppressive tumor microenvironment, and driving immunotherapy resistance.

**Highlights:** High FSTL3 levels are associated with poor outcomes in ovarian cancer.

Serum levels of FSTL3 increase during tumor growth and reflect tumor burden and therapy response.

Overexpression of FSTL3 in cancer cells promotes a fibrotic tumor microenvironment and immunocyte exclusion.

Overexpression of FSTL3 in tumors induces resistance to Chk1 and immune checkpoint inhibitor combination therapy.

**Graphical abstract:** 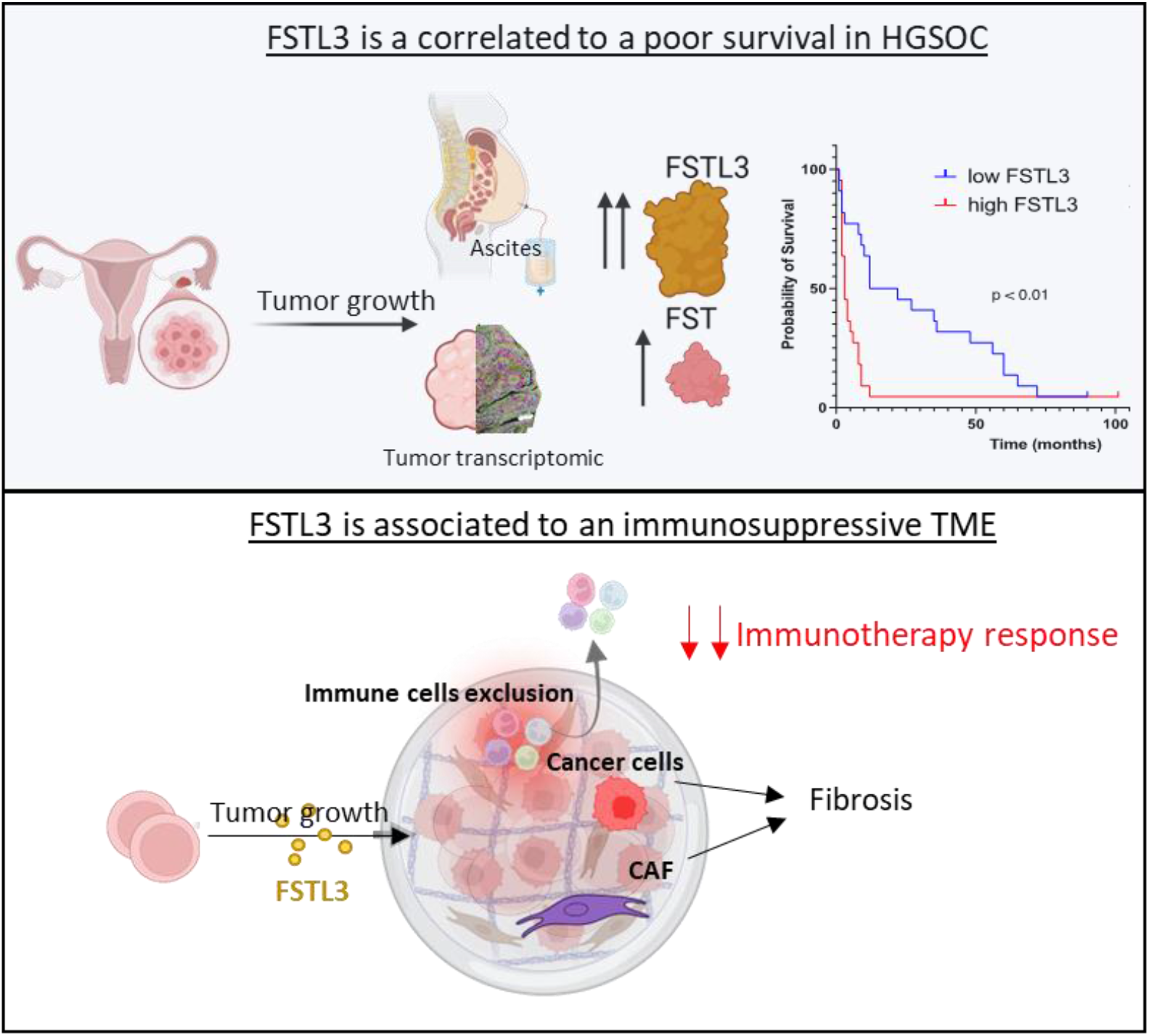

## Introduction

Ovarian Cancer is the 8th most common cancer among women worldwide (1, 2). with over 324,000 new cases and approximately 207,000 deaths reported in the world in 2022, making it the 5th leading cause of cancer-related mortality in women (1, 2). Projections indicate that by 2050, the worldwide incidence of ovarian cancer will increase by over 55%, and the number of deaths due to ovarian cancer is set to rise by almost 70%, to a total of 350,956 per year (1, 2). The high mortality rate is largely attributed to late-stage diagnosis, as the disease often presents with nonspecific symptoms. Ovarian carcinoma accounts for over 90% of all ovarian cancer cases, with high-grade serous ovarian carcinoma (HGSOC) being the most prevalent and aggressive subtype (3, 4). The standard treatment for ovarian cancer remains debulking surgery combined with platinum-based chemotherapy. Despite optimal treatment, prognosis remains poor, with 70-80% of patients experiencing relapse, and a 5-year survival rate of only 17% for those diagnosed at advanced stages, primarily due to resistance to platinum-based therapies (5, 6).

Platinum-resistant recurrent HGSOC, particularly in BRCA wild-type cases (which represent approximately 75% of all HGSOC cases poses a significant therapeutic challenge due to limited treatment options (7–9). This highlights the urgent need to develop novel therapeutic agents. Despite the high prevalence of CD4+ and CD8+ T cells, immune checkpoint blockade therapies, designed to enhance anti-tumor immune responses, have demonstrated limited efficacy in ovarian cancer(10–12). This may be attributed to the tumor microenvironment, which is adept at suppressing immune responses by recruiting and promoting immune-suppressive cells, thereby evading the effects of immunotherapy (13–15).

Recent advancements in preclinical models have provided valuable insights into treatment resistance. The KPCA syngeneic mouse model of CCNE1-amplified HGSOC bears **K**rasG12V and Tr**p**53R172H mutations and overexpresses **C**cne1 and **A**kt2 (16). We have previously observed significant variability in therapeutic responses among genetically identical cell line clones of KPCA treated with a combination of immunotherapy (anti-**P**D-L1/**C**TLA-4) and targeted therapies (**P**rexasertib, CHK1 inhibitor) (collectively referred to as PPC), ranging from clones with complete sensitivity (KPCA.B) to complete resistance (KPCA.C). When comparing the transcriptomes of these clones we identified follistatin (FST) as a potential immunomodulatory factor overexpressed in tumors resistant to PPC treatment (16). To further investigate the mechanisms underlying this resistance, the follistatin gene was deleted in KPCA.C resistant clone (KPCA.FSTKO), which restored sensitivity to PPC; likewise, overexpression of FST in the sensitive KPCA.B clone (KPCA.B-FST^OE^) made it resistant to PPC treatment (16). We hypothesized that follistatin inhibits immune responses, and thus promotes immunotherapy resistance, through the neutralization of TGF-β superfamily ligands, such as Activin A, which plays a critical role in recruiting immunocytes (17, 18). Elevated follistatin expression in tumors was associated with poorer overall survival in CCNE1-amplified HGSOC, and its presence in blood may serve as an early diagnostic marker for ovarian cancer(19).

Building on these findings, we sought to investigate the role of follistatin-like-3 (FSTL3), a close paralog of FST with overlapping TGF-β superfamily ligand affinity (20). A recent study in colorectal cancer suggested that FSTL3 contributes to immune evasion and showed that FSTL3 was overexpressed in malignant cells and was linked to poor clinical survival outcomes with anti-PD1 therapy (21). In immunocompetent tumor models, FSTL3 knockout led to an increased proportion of CD8+ T cells and a reduction in regulatory T cells and exhausted T cells, improving the efficacy of anti-PD1 therapy (21). This suggests that FSTL3 could serve as both a biomarker for immunotherapeutic efficacy and a novel therapeutic target to sensitize tumors to anti-PD1 treatment (21).

This study aimed to understand how FSTL3 contributes to ovarian tumor development and aggressivity, and if it regulates the tumor immune microenvironment. We explored the role of FSTL3 independently of FST by overexpressing human FSTL3 in the KPCA.FSTKO cell line, which we will refer to as KPCA.FSTKO_hFSTL3. In this study, we found that overexpression of FSTL3 changed the tumor microenvironment, leading to increased fibrosis and immune exclusion, and caused resistance to immunotherapy (PPC treatment).

## Results

### 1) Clinical Investigation of FST and FSTL3 in Ovarian Cancer: Insights from Patient Samples

**Figure 1:**
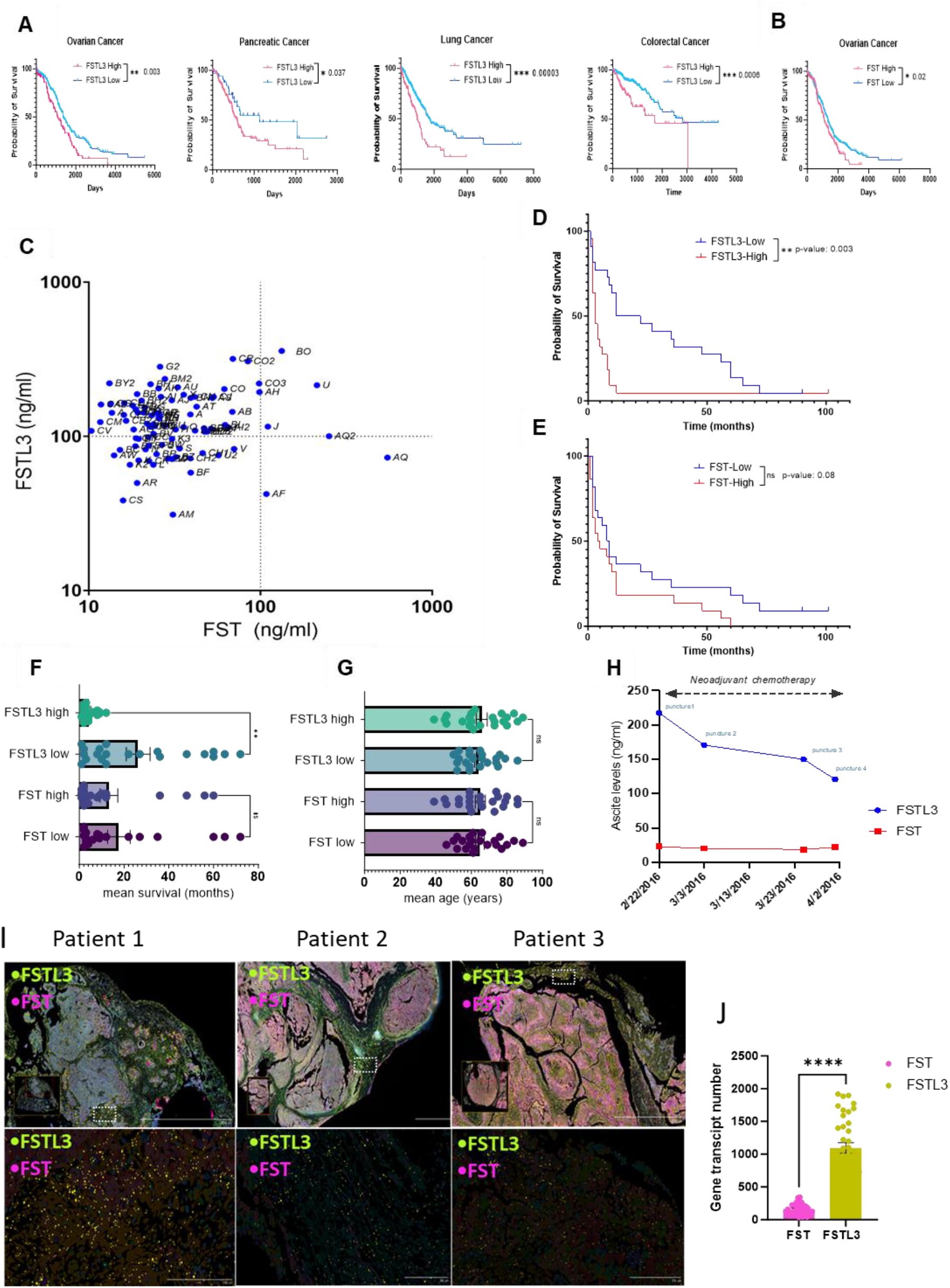
FSTL3 is overexpressed in ascites and human tumors section and is associated with poor survival. A) Kaplan-Meier survival plots comparing patient populations with low and high FSTL3 and B) FST transcriptomic expression according to the TCGA dataset. C) Concentration of FST and FSTL3 in 96 ascites samples from patients with various ovarian cancer subtypes. D-E) Kaplan-Meier survival curves comparing low and high FST and FSTL3 protein levels in 44 patients with HGSOC, BRCA1/2 mutation-negative (groups defined by median expression levels).F) Mean survival (in months) comparing patients with low and high FST and FSTL3 levels from panels D-E. G) Mean age distribution across groups from panel D-E. H) FST and FSTL3 protein levels measured in ascites during neoadjuvant chemotherapy in a 44-year-old patient with HGSC. I) Spatial transcriptomics was performed on HGSOC tissue samples from three patients using the Xenium (10X genomics). The images display a representative section, showing the spatial expression of FST and FSTL3 transcripts within the tissue microenvironment. J) Panel presenting the number of transcripts of FST and FSTL3 (N=3).

### 2) FST and FSTL3 are supraphysiological expressed in ascites of ovarian cancer patients

Analysis of The Cancer Genome Atlas dataset revealed that FSTL3 is associated with poor survival outcomes in multiple cancers, including ovarian, colorectal, pancreatic and lung cancers (Fig. 1A), whereas FST shows a weaker correlation of poor prognosis for ovarian cancer (Fig. 1B). To evaluate the relative expression of FST and FSTL3 in ovarian cancer, we measured the concentrations of these proteins by ELISA in 96 ascites samples from 77 patients with various types of epithelial ovarian cancer (mixed histologic types, mixed genotypes (TP53, BRCA1/2), both chemo-naïve and treated, and at different stages) (Table 1). We found that FST expression in ascites (mean: 42.5 ng/ml) was substantially elevated, ranging from 10- to 139-fold higher compared to normal peritoneal fluid (1.8 ng/mL) and blood serum (3.5 +/- 0.2 ng/ml) (22). Furthermore the FST levels in HGSOC ascites are higher than in the ascites of endometrioma patients (9.8ng/ml) (19, 23). Notably, FSTL3 levels in ascites (mean: 130.9ng/ml) were significantly higher than those of FST (p<0.0001), with a mean fold increase of 5.14 (Fig. 1C) (Table 1). Although normal FSTL3 concentrations in peritoneal fluid are not well documented, our findings indicate that these levels ranged from 10- to 100-fold higher than typical serum concentrations reported in the literature (mean: 3.4 ng/ml) (24).

### High FSTL3 levels in ascites are associated with poor survival in HGSOC

Next, we sought to examine the correlation between FST and FSTL3 levels in ascites and overall survival. To reduce variability and improve clinical relevance, we focused our analysis on a more homogenous group of 44 patients with high-grade serous ovarian cancer and BRCA wild-type genotype. We found that elevated FSTL3 levels were significantly associated with reduced survival rates (p=0.0035) (Fig. 1D). Notably, when dividing patients into two groups based on the median FSTL3 expression (median = 109.8 ng/mL, IQR [75.65–143.5]), the group with higher FSTL3 levels had a significantly shorter mean survival of 4.38 ± 0.67 months, compared to 26.33 ± 5.33 months in the group with lower FSTL3 levels (p<0.01) (Fig. 1F).

In contrast, although the correlation between FST levels and survival was less pronounced (p = 0.088) (Fig. 1E), we observed a mean survival of 17.7 ± 5.06 months in the group with lower FST expression (median = 26.00 ng/mL, IQR [21.35–38.21]), compared to 13.23 ± 3.96 months in the group with higher FST expression (Fig. 1F). However, this difference was not statistically significant. These findings were consistent with previous TGCA analysis (Fig. 1A/B). Importantly, no significant correlation was observed between patient age and the expression levels of either FST (mean: 42.5ng/ml) or FSTL3 (mean: 130.9ng/ml) (Fig. 1G, Table 1). Finally, when FST and FSTL3 levels were measured in ascites samples from a 44-year-old patient with HGSOC during her course of neoadjuvant chemotherapy (Carboplatin- Taxol), a reduction in FSTL3 but not FST levels was observed over time (Fig. 1H), suggesting FSTL3 concentration in ascites fluid may reflect tumor burden and response to therapy.

### FSTL3 is more highly expressed than FST in human HGSOC tumors

We aimed to investigate whether the expression levels of FST and FSTL3 in human ovarian tumor tissues were consistent with the patterns previously observed in ascitic fluid from ovarian cancer patients. To achieve this, we employed spatial transcriptomic analysis using the Xenium platform, a cutting-edge technology that enables high-resolution mapping of gene expression on tissue slides. Tissue sections were obtained from three treatment-naive high-grade serous ovarian cancer (HGSOC) tumors. Our analysis revealed that FSTL3 transcripts were expressed at significantly higher levels than FST, mirroring the trend observed in ascites samples (Fig. 1I). The quantification of gene transcripts was performed using the Xenium analysis software, which detects and counts individual transcripts. Transcript counts were averaged across 12 non-overlapping regions of interest per patient to ensure a comprehensive representation of all tumor areas and sample heterogeneity within the tumor microenvironment (Fig. 1J). Across the three patients analyzed, the mean transcript count for FSTL3 was 1094.64, which was significantly higher compared to FST, with a mean transcript count of 161.75 (Fig. 1J). These findings highlight the predominance of FSTL3 expression in both ascitic fluid and tumor tissues, suggesting a potential role for FSTL3 in the tumor microenvironment of HGSOC.

### 3) Serum FST and FSTL3 levels increase during tumor growth, and reflect tumor response to therapy, in a mouse model of ovarian cancer

#### Serum levels of FST and FSTL3 increase during tumor growth in mice

To determine if FST and FSTL3 serum levels correlate with tumor growth and can serve as biomarkers of tumor burden, we monitored their levels in a syngeneic mouse model of CCNE1-amplified high-grade serous carcinoma (KPCA.C cell line clone), which we have previously reported secretes FST (25). We measured the concentrations of FST and FSTL3 in the serum of mice implanted with the KPCA cancer cell line (23). Given that the ovaries are also expected to secrete high levels of FST (26), we compared intact mice to oophorectomized mice, to control for ovarian-dependent FST secretion, and measured serum levels of FST and FSTL3 by ELISA (Fig. 2). Blood samples were collected through submandibular vein puncture (cheek blood) and via cardiac puncture at the endpoint.

**Figure 2:**
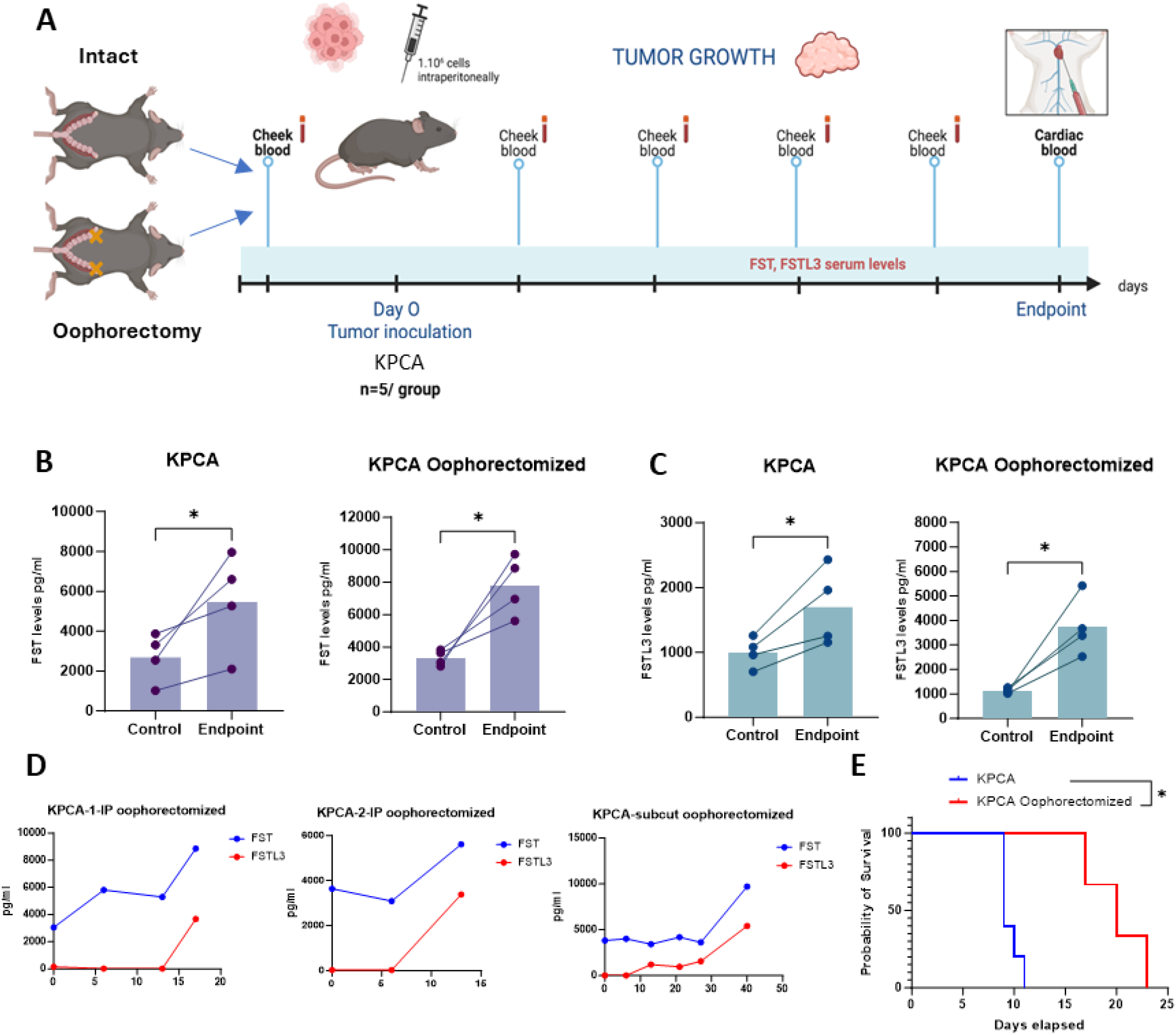
FST and FSTL3 serum levels correlate with tumor growth. A) In vivo experimental design outlining the timepoints of blood collection during tumor growth following implantation with KPCA.C cells in mice after oophorectomy, or in intact mice. B-C) Serum levels of FST and FSTL3 in intact and oophorectomized mice before grafting and at endpoint. D) FST and FSTL3 serum levels during tumor growth (1 & 2 KPCA injected IP in oophorectomized mice, 3-KPCA oophorectomized subcutaneous). E) Kaplan-Meier curves comparing KPCA graft model survival in intact and oophorectomized mice.

We observed a significant increase in FST and FSTL3 serum levels by ELISA at endpoint compared to pre-graft levels in the KPCA model (Fig. 2A-B) (N=4 mice per group). Similarly, in oophorectomized mice, FST and FSTL3 levels rose significantly during tumor growth. Surprisingly, oophorectomy in mice did not significantly alter the FST concentration between the pre-graft and endpoint measurements, suggesting there may be other confounding sources of FST. Longitudinal monitoring of FST and FSTL3 in serum (N=4), regularly tracked during tumor growth, revealed a similar upward trend for both proteins over time (Fig. 2D). Interestingly, ovariectomized mice grafted with both exhibited a significant prolongation of survival (p<0.05), suggesting an influence of ovarian hormonal in promoting tumor growth (Fig. 2E).

#### FSTL3 Overexpression induces Resistance to PPC treatment

To better understand the consequence of FSTL3 overexpression on ovarian tumor growth and response to therapy (Fig 3A), we used the parental cell line KPCA.FSTKO to generate a cell line overexpressing human FSTL3 (KPCA.FSTKO_hFSTL3). We developed this model both to allow us to interrogate the function of human FSTL3 and to enable the future development of specific inhibitors. The overexpression of FSTL3 in the resulting clone was validated by ELISA (Fig 3B).

**Figure 3:**
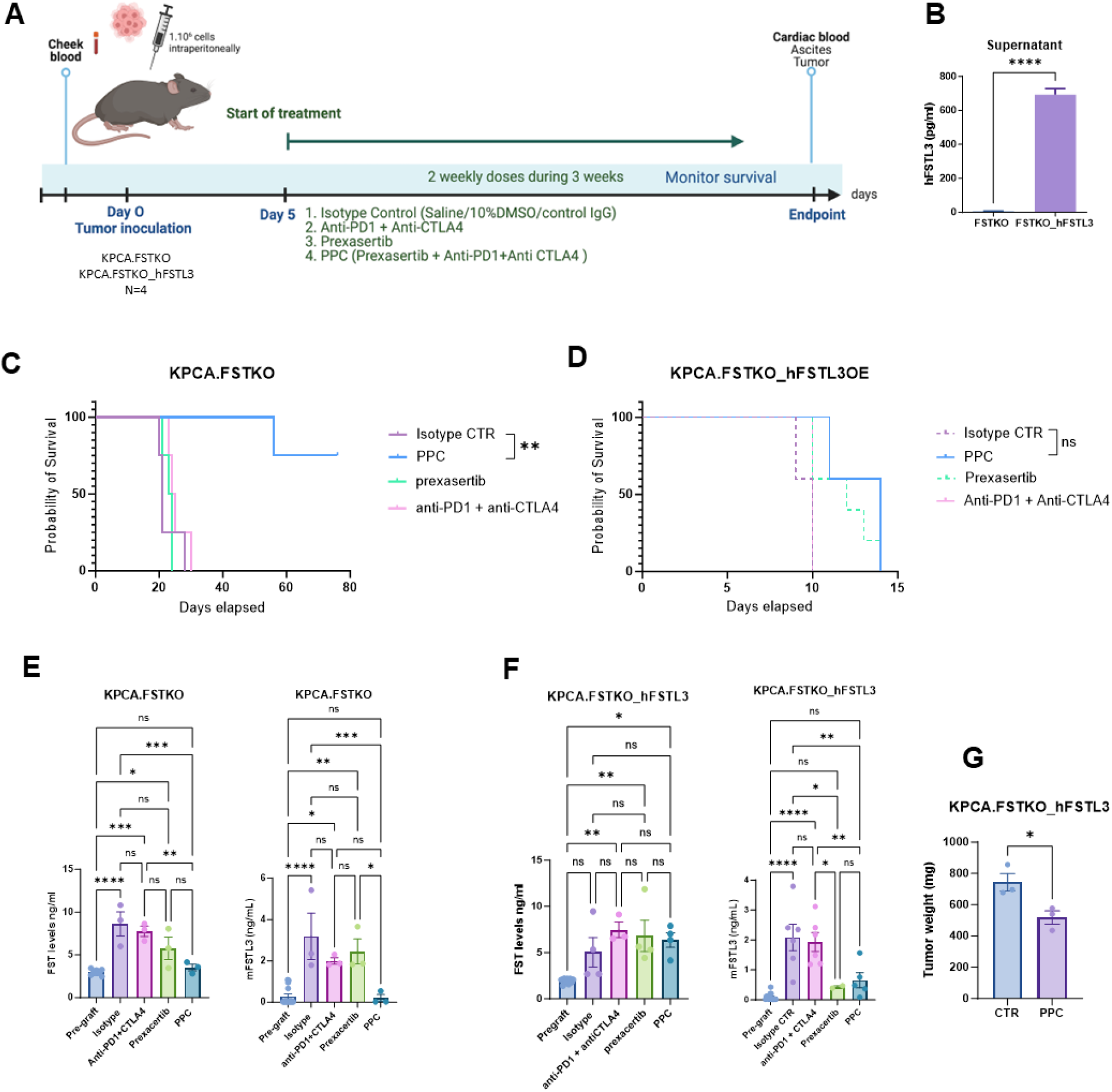
FSTL3 overexpression causes immunotherapy resistance. A) In vivo experimental design outlining the treatment groups and methodology used to evaluate PPC immunotherapy response. B) Validation of hFSTL3 overexpression by ELISA comparing the supernatant of media conditioned by the KPCA.FSTKO parental and chosen KPCA.FSTKO_hFSTL3 clone. C) Kaplan-Meier survival curves comparing the survival outcomes of KPCA.FSTKO and D) KPCA.FSTKO_hFSTL3 models following treatment with control (IgG isotype + vehicle), Prexasertib, anti-PD1 + anti-CTLA4, or the PPC triple combination (Prexasertib + anti-PD1 + anti-CTLA4) by intra-peritoneal injections 2 times/week (N=4/groups) E) Comparison of FST and FSTL3 serum levels pre-graft and at endpoint across all treatment groups in KPCA.FSTKO and F) KPCA.FSTKO_hFSTL3 grafted mice. G) Tumor weight at endpoint (day 10) in the KPCA.FSTKO_hFSTL3 model treated or not with PPC.

The KPCA.C syngeneic mouse model of HGSOC is inherently resistant to PPC treatment, which includes a combination of anti-PD1 (50ug), anti-CTLA4 (50ug), and the CHK1 inhibitor Prexasertib (10mg/kg), including two IP-injections per week (16). Previous findings indicated that the KPCA.FSTKO model, achieved through genetic ablation (KPCA.FSTKO), restored the sensitivity of KPCA.C to the PPC treatment (16). We hypothesized that FSTL3, being a close homolog of FST, could also induce resistance to PPC. As previously reported, we found that individual therapies such as Prexasertib, or a combination of immune checkpoint blockade alone (anti-PD1 plus anti-CTLA4) were not sufficient to prolong the survival of mice implanted with KPCA.FSTKO cells. However the combination of all three therapies (anti- PD1 plus anti-CTLA4 plus Prexasertib) or “PPC” was synergistic and significantly improved survival with 80% of the mice tumor-free at the end of the study (p<0.01) (Fig3C).

In contrast, overexpression of FSTL3 caused resistance to PPC in our KPCA.FSTKO_hFSTL3 model (Fig. 3D), indicating that overexpression of FSTL3 phenocopies FST-mediated resistance to PPC, which failed to significantly improve overall survival, albeit leading to slightly smaller tumors at endpoint (Fig. 3G).

#### PPC therapy restores physiological FST and FSTL3 Levels

We next sought to determine if the therapeutic response to PPC, and the corresponding decrease in tumor burden would be reflected in FST or FSTL3 levels in serum at endpoint. We observed significant increases in FST and FSTL3 levels when comparing pre-graft levels (FST 2.9 ng/ml, FSTL3 0.15 ng/ml) to endpoint levels following grafting with KPCA.FSTKO cells and treatment with either isotype + vehicle control (FST 8.6 ng/ml, FSTL3 3.1 ng/ml), anti-PD1 + anti-CTLA4 (FST 7.7 ng/ml; FSTL3 2 ng/ml), or Prexasertib (FST 5.7 ng/ml; FSTL3 2.45 ng/ml) alone, reflecting the muted effect of these treatments on survival. In contrast, PPC treatment significantly reduced FST and FSTL3 levels compared to isotype control and restored both FST (∼3.5ng/ml) and FSTL3 (∼0.19ng/ml) to physiological levels (Fig. 3E).

In the mice grafted with KPCA.FSTKO_hFSTL3 cells, none of the treatments caused significant changes in the FST levels, suggesting overexpression of hFSTL3 induced resistance to PPC treatment in these tumors (Fig. 3F). Interestingly, when measuring endogenous mouse FSTL3 levels, we observed a significant reduction in concentration compared to control (Fig. 3F), suggesting FSTL3 may be a more specific biomarker of tumor burden. This decline in FSTL3 may be due to a modest effect of PPC treatment on tumor burden reduction (fold change: 1.31), as observed at endpoint (p < 0.05) (Fig. 3G).

### 4) FSTL3 overexpression promotes tumor growth and a more mesenchymal tumor microenvironment

#### FSTL3 promotes Tumor growth and is associated with a more fibrotic microenvironment

To investigate the consequences of FSTL3 on tumor growth, we compared tumor growth in established models (KPCA.FSTKO and KPCA.FSTKO_hFSTL3). Following implantation intraperitoneally at 1.10^6 cells/mouse, we observed that FSTL3 overexpression led to a 50% higher tumor mass in mice receiving KPCA.FSTKO_hFSTL3 compared to the KPCA.FSTKO cells at day 10 post-graft (Fig. 4A). Additionally, survival analysis revealed a significant decrease in the survival of mice bearing KPCA.FSTKO_hFSTL3 derived tumors compared to those implanted with KPCA.FSTKO (Fig. 4B).

**Figure 4:**
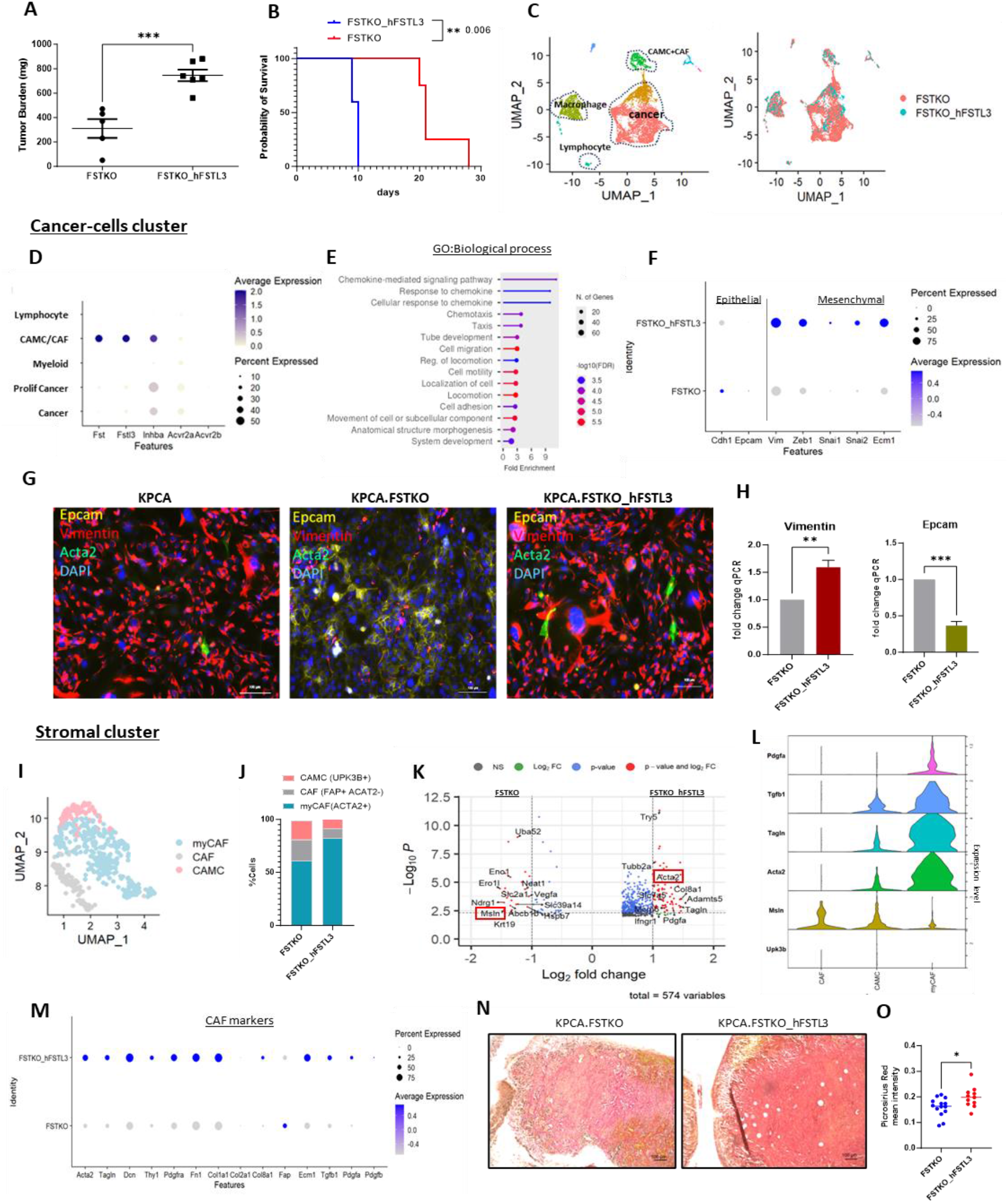
Overexpression of FSTL3 increase tumor development and is associated to fibrotic microenvironment. A) Tumor mass of KPCA.FSTKO and KPCA.FSTKO_hFSTL3 model harvests from peritoneal cavity of mice at endpoint. B) Kaplan Meier of the survival of KPCA.FSTKO and KPCA.FSTKO_hFSTL3. C) UMAP of cell population from the scRNA seq of KPCA.FSTKO and KPCA.FSTKO_hFSTL3. D) DotPlot representing the transcript expression levels of Fst, Fstl3, Inhba, Acvr2a, and Acvr2b across different cell identities within the tumor microenvironment of KPCA.FSTKO and KPCA.FSTKO_hFSTL3 models from scRNAseq. Dot size corresponds to the percentage of cells expressing the gene. Dot color intensity indicates average expression level, with darker colors representing higher expression. E) Pathway analysis (GO biological process) of differential gene expression of KPCA.FSTKO and KPCA.FSTKO_hFSTL3 in cancer cluster. F) Dot plot illustrates the expression patterns of key EMT-related genes in cancer cells cluster from KPCA.FSTKO and KPCA.FSTKO_hFSTL3. G) Representative images show the expression of Epcam (epithelial marker, yellow), Vimentin (mesenchymal marker, red), Acta2 (myofibroblast marker, green), and nuclei stained with DAPI (blue) by immunofluorescence. H) RT-qPCR of Epcam and Vimentin expression of both KPCA.FSTKO and KPCA.FSTKO_hFSTL3. I) UMAP of cluster showing the diverse subclusters of stromal compartment including Cancer Associated Mesothelial Cells (CAMC), Cancer Associated Fibroblast (CAF) and myofibroblast (myCAF). J) Proportion (% cells) for each subcluster (CAMC, CAF, myCAF) in KPCA.FSTKO and KPCA.FSTKO_hFSTL3 of TME. K) Volcano Plot of differential gene expression of in KPCA.FSTKO and KPCA.FSTKO_hFSTL3 in stromal cluster. L) VlnPlot showing the expression of secreted factors (TGFB1 & PDGFA) in CAMC (UPK3K+, MSLN+) and myCAF subtypes (ACTA2+, TAGLN+). M) DotPlot representing a panel of CAF markers including collagen, extracellular matrix organization and secreted factors transcripts. N) Image representative tumor section of KPCA.FSTKO and KPCA.FSTKO_hFSTL3 stained with Sirius Red. O) Quantification of red Sirius staining N=10)

To investigate the consequence of FSTL3 overexpression on the composition of the TME we conducted an scRNAseq experiment in the KPCA.FSTKO and KPCA.FSTKO_hFSTL3 tumor models (day 10 post- graft) (Fig. 4C). We hypothesized that FSTL3 overexpression in cancer cells not only modifies the cancer cell phenotype, but also modulates immunotherapy response through its action on stromal cell types of the tumor microenvironment. Thanks to the use of a human *hFSTL3* transgene in the KPCA.FSTKO_hFSTL3 model, we were able to mask the exogenous human FSTL3 and specifically detect endogenous murine sources of FSTL3 in the tumor microenvironment by scRNAseq, and by murine- specific ELISA. Transcriptomic analysis of KPCA.FSTKO and KPCA.FSTKO_hFSTL3-derived tumor revealed that endogenous mouse FST and FSTL3 were predominantly secreted by cancer-associated mesothelial and fibroblast cells, which are major components of the stromal tumor microenvironment (Fig. 4D). In contrast, activin A (*Inhba*), one of the main ligands bound by both FST and FSTL3 was primarily secreted by cancer cells, while its type II receptors (*Acvr2a, Acvr2b*) were highly expressed in both cancer cells and fibroblasts (Log2FC > 0.5) (Fig. 4D).

When we compared the differential gene expression in the cancer clusters from KPCA.FSTKO and hFSTL3-overexpressing tumors, we found a significant enrichment in differentially expressed genes related to pathways in cell migration, adhesion, and locomotion (GO biological process) (Fig. 4E). Underlying these processes, cancer cells overexpressing FSTL3 exhibited increased expression of the most frequent genes associated with epithelial-mesenchymal transition (*Vim*, *Zeb-1*, *Snai1/2*) and extracellular matrix organization (ECM1), while expression of epithelial markers (Epcam, Cdh1) decreased (Fig. 4F) (27, 28). The differential expression of epithelial (Epcam) and mesenchymal (Vimentin) markers was confirmed by immunofluorescence and qPCR (Fig. 4G/H).

We next evaluated how FSTL3 overexpression by cancer cells altered the stromal cells in the TME, including cancer-associated mesothelial cells (CAMC) and cancer-associated fibroblast (CAF). Firstly, among stromal cell types, three subtype clusters were identified based on marker expression including CAMCs (*Msln+*), CAFs (*Fap*+), and MyCAFs (myofibroblasts, *Acta2*+) (Fig. 4I). The proportion (percentage of cell) of CAF, MyCAF, and CAMC cells in the stromal cluster was quantified in each model. The KPCA.FSTKO_hFSTL3-derived tumors had a higher proportion of myCAFs (82%) and a lower proportion of CAMC cells (9%) than KPCA.FSTKO tumor with 61% myCAFs and 18% CAMCs in the stromal compartment (Fig. 4J). We next compared the differential gene expression between KPCA.FSTKO and KPCA.FSTKO_hFSTL3 stromal cluster and observed a significantly increased expression (Log2FC>0.5) of myCAF markers (*Acta2, Tagln*), collagen (*Col8a1)*, and secreted factor (Pdgfa) consistent with a myofibroblast phenotype in the KPCA.FSTKO_hFSTL3 tumor. In contrast, in the KPCA.FSTKO tumors, the stromal cluster expressed higher levels of mesothelial markers such as Krt19 and Msln, consistent with a higher proportion of CAMC in this model (Fig. 4K). In addition, we revealed that Tgfb1 was more highly expressed by myCAF compared to CAMC and CAF cells (Fig. 4L). Since myofibroblast can promote fibrosis and immune evasion (29–33), we next evaluated whether FSTL3 overexpression altered these processes. We compared the expression of collagen, fibronectin, and other extracellular matrix markers that contribute to fibrosis. We found that collagens (*Col1a1*, *Col2a1*, *Col8a1*), fibronectin (*Fn1*), and extracellular matrix component (*Ecm1*) were increased when FSTL3 was overexpressed in TME (Fig. 4M). Furthermore, the secreted factors *Tgfb1* and *Pdgfa*, which are known to promote fibrosis (34), were also more highly expressed in KPCA.FSTKO_hFSTL3 tumors (Fig. 4M). In addition, we confirmed that the tumors overexpressing FSTL3 exhibited a significantly higher collagen deposition within the tumor microenvironment as represented by the Sirius Red staining (Fig. 4N/O), suggesting increased fibrotic areas within these tumors (35).

#### FSTL3 overexpression in TME promotes immune exclusion

Given the immunosuppressive role of myCAFs and fibrosis, we investigated whether FSTL3 overexpression affected the infiltration of immune cells, and particularly T cells, given their requirement for PPC treatment response (16). To this end, we dissociated three tumors per group from treatment-naïve KPCA.FSTKO and KPCA.FSTO_hFSTL3 bearing mice. We stained the cell suspension with specific antibodies, enabling us to quantify immune cell populations by flow cytometry. We measured the total percentage of immune cells (CD45+) and T-cell subsets (CD4+, CD8+) present in the TME. We observed a significant reduction in overall immune cell infiltration, with an average decrease of 50% (CD45+) in tumors overexpressing FSTL3 (Fig. 5A/B). In addition, T cells were markedly reduced in FSTL3-overexpressing tumors compared to controls with CD4+ (25% versus 15%) and CD8+ (15% versus 5%), suggesting FSTL3-induced immune exclusion may participate in PPC treatment resistance (Fig. 5C).

**Figure 5:**
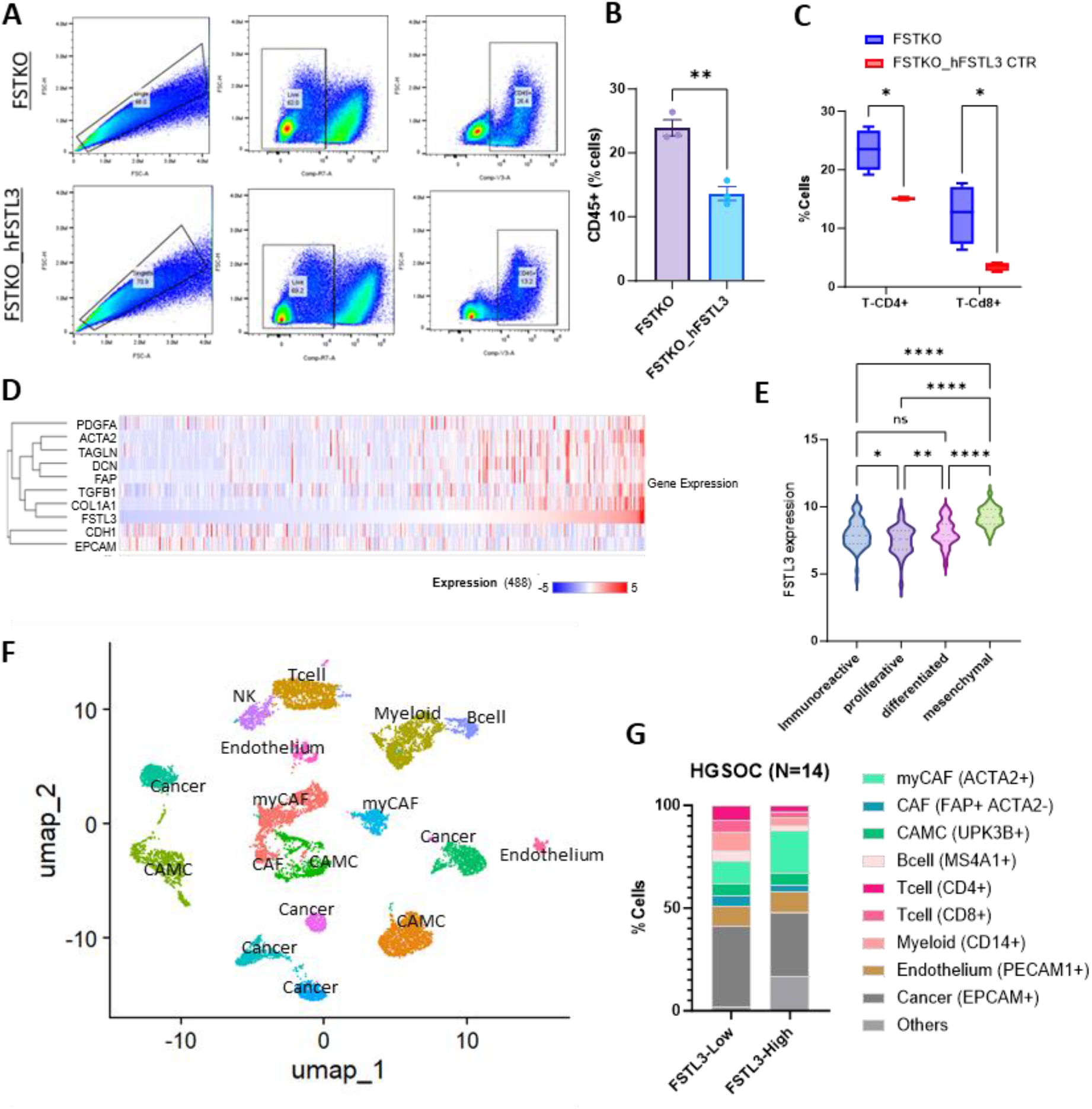
FSTL3 overexpression promotes immunocyte exclusion. A) Gating strategy to isolate singlet, live, CD45+ cells from KPC.FSTKO and KPCA.FSTKO_hFSTL3 tumors. B) Percent of CD45 positive cells in KPCA.FSTKO and KPCA.FSTKO_hFSTL3 tumors (N=3). C) Percent of T cells (CD4/CD8,) gated on CD45 positive live cells in KPCA.FSTKO and KPCA.FSTKO_hFSTL3 tumors (N=3). D) Heatmap of relative transcript expression of selected genes correlated to FSTL3 expression across patient samples from the TCGA (HGSOC, 486 cases), with red indicating higher expression and blue indicating lower expression. E) Analysis of TCGA HGSOC tumor transcriptomes (N=308) showing expression of FSTL3 by molecular histotype. F) UMAP of HGSOC of 14 patients from dataset:GSE235931. G) Proportion of cells present in TME between patients with high and low FSTL3 expression based on median expression (N=7 per group).

To determine if FSTL3 overexpression is also associated with a more fibrotic and immunosuppressive TME in human HGSOC tumors, we data from The Cancer Genome Atlas (TCGA) dataset of bulk RNAseq from 486 HGSOC cases (36). We found an association between high FSTL3 transcript levels and expression of CAF markers and fibrosis markers (COL1A1, TGFB1, FAP, DCN, TAGLN, ACTA2, PDGFA), previously identified as upregulated in KPCA.FSTKO_hFSTL3 mouse tumors. In contrast, epithelial markers (EPCAM, CDH1) showed an inverse correlation with FSTL3. We next sought to evaluate if FSTL3 overexpression was associated with a particular ovarian cancer molecular histotype (7). To do so we compared FSTL3 transcript levels from an RNAseq dataset of the TCGA ovarian cancer UNC hub (n=308) annotated with the molecular subtypes as proliferative, immunoreactive, differentiated, and mesenchymal, and found that FSTL3 was significantly higher in the mesenchymal subtype compared to all others (Fig3N). These data suggest that FSTL3 overexpression also promotes a more mesenchymal phenotype in human tumors.

To further confirm the role of FSTL3 in inducing immune evasion in human patients, we analyzed the TME using single-cell RNA-seq data from fourteen HGSOC patients. Patients were stratified into two groups based on median FSTL3 expression levels: FSTL3-high (N=7) and FSTL3-low (N=7). We then compared the cellular composition of the TMEs between the two groups based on % marker expression. Similar to observations in the mouse TME, FSTL3-high human samples exhibited an increased proportion of stromal cells (myCAF and CAMC) and a marked reduction CD4+and CD8+ T cells respectively. These findings suggest that FSTL3 overexpression also promotes a high abundance of CAFs and T cell exclusion in human tumors.

## DISCUSSION

Ovarian cancer, and particularly HGSOC, continues to pose a significant clinical challenge due to its frequent late-stage diagnosis and high recurrence rates. The identification of FST as a potential player in immune evasion and treatment resistance has paved the way for exploring the role of other follistatin homologs such as FSTL3 in ovarian cancer (16).

Notably, this study identified significant overexpression of both FST and FSTL3 in 96 ascites samples from ovarian cancer patients. In particular, we found that FSTL3 levels were generally higher than FST, both in the ascites and in the tumors of HGSOC patients. Moreover, elevated FSTL3 protein levels in ascites of patients with BRCA1 wildtype tumors, where treatment options are notably limited, were significantly associated with poorer survival outcomes. This correlation between overexpression of FSTL3 and poor outcome may not be unique to ascites nor ovarian cancer as it was also observed at the transcriptional level across multiple solid tumor types including pancreatic, lung, colorectal and ovarian through analysis of data from the TCGA. These findings emphasize a potential role for FSTL3 as a prognostic biomarker for disease progression.

Importantly, we observed a strong relationship between FSTL3 expression in the serum and tumor growth, and marked decreases in FSTL3 associated with treatment responses in mice. Similarly, FSTL3 levels decreased during neoadjuvant chemotherapy in a patient with serial sampling of ascites. These data suggest FSTL3 may have clinical utility in monitoring tumor burden and treatment efficacy. Importantly, FSTL3 background levels in the blood of mice were low, and FSTL3 levels in humans were not associated with patient age, reinforcing its broad potential as a biomarker to track HGSOC progression.

In a previous study we showed that knocking out FST in the “KPCA” model of CCNE1-amplified HGSOC promoted sensitivity to a combination immunotherapy treatment (PPC: Prexasertib, PD- 1, and CTLA-4 blockade) (16). In this study, we show that overexpression of human FSTL3 in this model (KPCA.FSTKO_hFSTL3) restores complete resistance to PPC, highlighting the redundant function of FST and FSTL3 in mediating immunotherapy resistance. The effectiveness of PPC therapy in the KPCA.FSTKO model was found to result from a synergistic effect between Prexasertib and immune checkpoint inhibitors (anti-PD1 and anti-CTLA4). Prexasertib, a checkpoint kinase 1 (CHK1) inhibitor, has shown potential for cancers that exhibit high DNA replication stress and impaired DNA damage response pathways (37, 38), such as HGSOC. A phase II trial studying Prexasertib in BRCA wild-type, platinum-resistant HGSOC showed some tumor shrinkage but limited overall responses (39), highlighting the clinical need to uncover combination therapies that can potentiate targeted therapies and overcome immunotherapy resistance. Indeed, immune checkpoint blockade has been largely ineffective in ovarian cancer, and the mechanisms of resistance remain elusive (40).

We hypothesized that FSTL3 contributes to immunotherapy resistance by promoting a mesenchymal tumor phenotype, increasing fibrosis by promoting myCAF development, and excluding immunocyte infiltration, particularly T cells. Additionally, FSTL3 may induce the secretion of other factors such as PDGF (Platelet-Derived Growth Factor) and TGF-β1 (Transforming Growth Factor Beta 1), which in turn promote fibroblast growth, migration, and differentiation (41, 42), further contributing to an immune-suppressive TME. Aberrant PDGF signaling has been implicated in the activation of stromal cells, such as fibroblasts, which promote an immunosuppressive niche (43). Similarly, Elevated TGF-β1 levels in the tumor microenvironment drive epithelial-to-mesenchymal transition (EMT), immune evasion, and extracellular matrix remodeling, processes that are critical for tumor invasion and metastasis (44). Moreover, TGF-β1 is a potent immunosuppressive cytokine that inhibits T-cell function and supports regulatory T-cell differentiation, thereby undermining anti-tumor immunity (45, 46). CAFs and other stromal cells such as CAMCs promote immune evasion (47–49) and may limit the effectiveness of immunotherapy treatments in solid tumors (28, 50). Fibrotic processes are common in cancers such as pancreatic, breast, lung, and liver cancer, where extensive fibrosis is frequently linked to poor prognosis (51) and can reduce immunocyte infiltration (32, 34, 52). In our mouse model, FSTL3 overexpression led to increased collagen deposition, and a 50% reduction in immune cell infiltration compared to controls. Notably, the populations of CD4+ and CD8+ T cells were markedly reduced in the FSTL3-overexpressing tumors in both mouse and human models. A similar immune profile was also observed in lung adenocarcinoma, where high FSTL3 levels correlated with a reduction of immune cell infiltration, including T lymphocytes and B lymphocytes in the TME (53). Our observations in an ovarian syngeneic tumor model largely parallel other studies in gastric and colorectal cancer where high FSTL3 was similarly associated with poor prognosis, EMT, fibrosis, and changes in immunocyte composition; however, the precise mechanism leading to these changes, and particularly the ligands modulated by FSTL3 in the context of the TME remain to be identified in ovarian cancer (54–58).

In summary, our data suggest that ovarian tumors secrete FSTL3 in the blood and ascites and that it is an indicator of poor prognosis. Experimental overexpression of FSTL3 in murine ovarian cancer changed the TME by promoting cancer cell EMT, increasing stromal fibrosis, and reducing immunocyte recruitment, which we speculate contributed to the induced resistance to immunotherapy treatment. Thus, FSTL3 may represent both a new secreted biomarker to optimize patient stratification, monitor disease burden and treatment response, and a new therapeutic target for combination immunotherapies in ovarian cancer. Further research is needed to build upon these findings and develop strategies to neutralize FSTL3 to meet the urgent demand for more effective treatments in HGSOC.

## Materials and Methods

### Human samples

Ascites samples from patients with various subtypes of ovarian cancers (96 samples) along with sections of high-grade serous ovarian cancer tumors were generously donated by female patients to the Massachusetts General Hospital, in accordance with an IRB-approved protocol (2007P001918).

Tumor sections used for spatial transcriptomics were donated by the Biological Resources Center of Montpellier Cancer Institute (ICM), France, and were collected following the French regulations under the supervision of an investigator. The collection was declared to the French Ministry of Higher Education and Research (declaration number DC-2008–695).

### ELISA

ELISA assays for human samples were performed using ANSH lab FST and FSTL3 kits. For mouse serum and ascites samples, RayBio mFST and AVIVA mFSTL3 kits were employed. ELISAs were conducted according to the manufacturer’s protocols. Optimal dilutions were determined before processing all samples. Absorbance was measured at 450 nm using a microplate reader (Pherastar), and protein concentrations were quantified based on standard curves generated following the kit instructions.

### Cell lines and 2D cell culture

Ovarian cancer cell lines KPCA, KPCA.FSTKO, KPCA.FSTKO_hFSTL3 were maintained in Dulbecco’s modified Eagle’s medium tissue culture medium supplemented with 5% fetal bovine serum (FBS) and 1% penicillin-streptomycin. All cells were cultured at 37°C in 5% CO2.

To generate the KPCA.FSTKO_hFSTL3 cell line we used the previously described KPCA.FSTKO cell line (16), and transfected it with a pcdna3.1 expression plasmid for hFSTL3 (Genscript). A stable clone was selected under geneticin selection based on FSTL3 secretion in conditioned media as measured by ELISA.

### RNA isolation and quantitative real-time PCR (RT-qPCR) analysis

Total RNA was isolated from cancer cells or tumors using the Qiagen RNA extraction kit. The cDNA synthesis was carried out with the SuperScript III First-Strand Synthesis System for RT-PCR (Invitrogen). The cDNA was then combined with primers and iQ SYBR Green Supermix (#1708882, Bio-Rad) in a 96- well plate. Reverse transcription was performed on a T100 Thermal Cycler (Bio-Rad), and qPCR was conducted using the CFX96 Touch Real-Time PCR Detection System (Bio-Rad). Gene expression levels were determined relative to housekeeping genes (GAPDH or HPRT) pre-designed from IDT biotech©, with expression quantification based on cycle threshold (Ct) values transformed using the 2^(-ΔCt) method, and the results were analyzed and plotted with GraphPad PRISM.

### Single-cell RNA sequencing

Single-cell RNA sequencing was performed using a 10X Genomics 3’ kit following manufacturer’s instructions. We dissociated tumors derived from KPCA.FSTKO and KPCA.FSTKO_hFSTL3 grafted mice (N = 4 pooled tumors per genotype) as previously described (47). We targeted 10,000 cells per sample, and obtained a total final cell count of 7582 for KPCA.FSKO and 4244 for KPCA.FSTKO_hFSTL3 after data analysis and quality control. The data analysis was performed using Cell Ranger (version 3.1.0) with standard parameters. Samples were aligned against the refdata-gex-mm10-2020-A reference sequences and were analyzed using Seurat library (R version 4.3.3). Standard pre-processing workflow was applied independently to each sample. Samples were filtered for mitochondrial percentage <20% and unique feature counts over 200 and less than 8000. Samples were normalized then merged using ‘‘merge’’ function in Seurat. Standard parameters for visualization and clustering were performed throughout the analysis. Markers for each level of cluster were identified using FindAllMarkers in Seurat.

The 10X scRNAseq datasets are available on the Gene Expression Omnibus (GEO) platform under accession number GEO: GSE283618

### Spatial transcriptomic

The spatial transcriptomic analysis was conducted using the Xenium platform (10x Genomics) on three treatment-naive high-grade serous ovarian cancer tumor samples. The expression levels of FST and FSTL3 transcripts were quantified across 12 representative regions per tumor to capture the spatial heterogeneity. The data was processed and analyzed using Xenium software (10x Genomics), enabling precise transcript quantification and spatial mapping of gene expression patterns.

### Single-cell analysis of patient datasets

FSTL3 transcript expression data were extracted from the GSE235931 dataset of 14 high-grade serous ovarian cancer tumors and analyzed using RStudio with Seurat. Patients were categorized into two groups based on their FSTL3 expression levels: FSTL3-High patients had an average expression above the population median (≥0.08), while FSTL3-Low patients had expression levels below the population median (<0.08). The tumor microenvironments of the FSTL3-High and FSTL3-Low groups were compared using specific markers indicative of cell phenotypes of interest, including ACTA2, FAP, UPK3B, CD4, CD8, EPCAM, MS4A1, and CD14. To further analyze the differences between these groups, the percentage of cells expressing these specific markers was calculated using the WhichCells command from the Seurat package.

### TCGA transcriptomic analysis

Survival analysis based on FSTL3 and FST expression was conducted using data extracted from The Cancer Genome Atlas (TCGA) from 349 ovarian cancer patients. Additionally, FSTL3 expression was analyzed in relation to survival outcomes in 497 lung cancer patients, 176 pancreatic cancer patients, and 307 colorectal cancer patients (Ov-TCGA, LUSC-TCGA, PAAD-TCGA, COAD-TCGA datasets)(59, 60). Patients were categorized into FST/FSTL3-High or FST/FSTL3-Low groups based on their expression levels relative to the median expression across all patients. Specifically, FST/FSTL3-High was defined as expression above the median, while FST/FSTL3-Low was defined as expression below the median. Statistical analyses were performed using GraphPad Prism, with survival curves compared using the Mantel-Cox log-rank test.

To evaluate FSTL3 expression in HGSOC by molecular histotype, we extracted FSTL3 transcript levels from an RNAseq dataset of the TCGA ovarian cancer (OV) UNC hub subset (n=308) with metadata identifying gene_expression_subtypes as proliferative, immunoreactive, differentiated, and mesenchymal. This level 3 data was downloaded from the UCSC Xena repository (61). The transcript numbers from Illumina HiSeqv2 data were log-transformed (log2(norm_count+1).

### Sirius red staining

Paraffin-embedded tissue sections were deparaffinized and rehydrated through graded alcohols to water. The sections were stained in a picro-Sirius Red solution, prepared by dissolving 0.5 g of Sirius Red (Sigma Aldrich, #365548) in 500 mL of a saturated aqueous solution of picric acid, for one hour. After staining, the sections were washed in two changes of 0.1M glacial acetic acid. Excess water was removed from the slides by blotting with damp filter paper. The slides were then dehydrated in three changes of absolute ethanol, cleared in xylene, and mounted in a resinous medium (VectorLabs). Picrosirius Red intensity was analyzed using the CellProfiler Software. Briefly, following image acquisition by light microscopy, the red channel marking collagen and yellow channels were split and the yellow was subtracted from the red one. The "smooth" module was then used before identifying the area covered by the stain. The mean intensity of this delimitated area was then quantified.

### Flow cytometry analysis

After dissociation of KPCA.FSTKO (N=4) and KPCA.FSTKO_hFSTL3 tumors (N=4), 5×10^4 cells were added in each well of a 96-well V-bottom plate. Cells were then immunostained with antibodies according to the manufacturer’s recommendations, washed 3 times and fixed with 1% Paraformaldehyde. Live and dead reagent (Thermofisher, L10119) was used to discriminate the live cells and antibody panels were used for T cells characterization including panel 1: [CD45+; CD8+] and panel 2: [CD45+; CD4+]. Immunostained cells were run on Cytek Aurora Flow cytometer and analyses were performed with FlowJo software.

### Immunofluorescence

Tumors were fixed overnight in 10% neutral buffered formalin. The fixed samples were subsequently processed through an alcohol series, cleared with xylene, and embedded in paraffin. Sections were cut to a thickness of 10 µm and mounted onto glass slides. Dewaxing was performed by incubating at 60°C for 60 minutes, followed by rehydration through a series of graded ethanol washes. Antigen retrieval was achieved using citrate buffer in a pressure cooker. Slides were then incubated for 1 hour at room temperature in blocking buffer, followed by three washes with PBS. The primary antibodies (anti-Epcam, anti-Vimentin, anti-Acta2) used at the concentration recommended by the manufacturer (Thermofisher), were applied overnight at 4°C. After washing, the slides were incubated for 1 hour with a fluorophore- conjugated secondary antibody (anti-Mouse-488nm/ anti-rabbit-555nm, Thermofisher). To visualize the stained proteins, sections were counterstained with DAPI for 5 minutes and then washed with PBS. Coverslips were mounted using Vectashield mounting medium to prevent fluorescence quenching. Fluorescent signals were visualized and analyzed using an epifluorescence microscope.

### Animal Experiments

This study was performed according to experimental protocols 2009N000033 and 2019N000068, approved by the Massachusetts General Hospital Institutional Animal Care and Use Committee. All experiments were made with 12-16 weeks female C57BL/6J mice purchased from the Jackson Laboratory. In each experiment, we used 4 to 5 mice per group.

**Cancer cell graft:** In all in vivo experiments, 1.10^6^ cells of KPCA, KPCA.FSTKO, or KPCA.FSTKO_hFSTL3 were injected intraperitoneally.

**Blood samples:** were collected using a 5 mm lancet on the submandibular vein, alternating the collection site between cheeks. A one-week healing interval was observed between procedures, while ensuring compliance with the maximum blood volume limits. Upon euthanasia, cardiac blood collection was performed. Serum was separated by allowing the collected blood (submandibular vein or cardiac collection) to clot at room temperature for 30 minutes, followed by centrifugation at 2,000 x g for 10 minutes at 4°C. The supernatant (serum) was carefully collected, labeled and stored at -20°C.

**Treatments:** began on day 5 post-graft and were administered intraperitoneally twice a week with the indicated doses during 4 weeks for KPCA.FSTKO and 2 weeks for KPCA.FSTKO_hFSTL3. For immunotherapy, anti-CTLA4 (50 μg/mouse; Bio X Cell, BE0164) and anti-PD1 (50 μg/mouse; Bio X Cell, BP0273) antibodies were used. Prexasertib (Selleckchem, S7178; 10 mg/kg) was administered either as a monotherapy or in combination with the immunotherapies and was resuspended according to the manufacturer’s instructions. Control mice were injected with 10% DMSO (vehicle) and an isotype control antibody (InVivoMAb mouse IgG2a isotype control, unknown specificity & InVivoPlus rat IgG2a isotype control, anti-trinitrophenol).

**Bilateral oophorectomy** was performed under Isoflurane anesthesia. Aseptic precautions were followed, and all surgical instruments were autoclaved prior to the procedure. The animal preparation included shaving the fur around the incision site, placing the animal on a warming pad, applying surgical soap scrub three times, and draping the animal. Eye lubricant was applied to prevent dryness. Carprofen (2-5 mg/kg) was administered subcutaneously just before surgery and continued every 12-24 hours for up to 72 hours postoperatively. Small incisions were made in the dorsolumbar region through the skin and body wall. The ovaries were exteriorized, followed by section and hemostasis using a cauterizer. The fascia was closed with sutures, and the skin was closed with surgical clips. Triple Antibiotic ointment (Bacitracin 400 U, Neomycin 3.5 mg, Polymyxin B 5,000 U.) was applied once. Mice were closely monitored on a warming mat post-surgery and then transferred to a cage with warm bedding. They were continuously monitored until they were fully ambulatory. Staples were removed at day 7 under isoflurane anesthesia.

**Monitoring**: Mice were monitored regularly for health and survival according to an IACUC approved protocol, euthanasia was performed if animals exhibited signs of poor body condition or distress.

### Statistical analysis

Statistical analysis was performed using GraphPad Prism 10.1.1 software. An independent sample t-test was used unless otherwise specified. Paired t-test was used to compare baseline and endpoint FST or FSTL3 levels by mouse. For comparisons involving more than two groups, one-way ANOVA was employed, or Kruskal-Wallis was used when data distribution was non-Gaussian. A p-value of ≤ 0.05 was considered statistically significant. Data are represented as the mean ± standard error of the mean (SEM).

## Supporting information

Concentration of FST and FSTL3 in Ascites Samples from Patients and their Clinical Characteristics.

## Acknowledgments

We would like to thank Robert Weinberg and Patricia K Donahoe for their comments. This work was supported by the Koch-MIT Cancer Center Bridge Award (DP and Robert Weinberg), the ECOR-FMD fellowship (MC) and the Innovation Development Grant (DP) from the Massachusetts General Hospital, The TEAL award from the Department of Defense-Congressionally Directed Medical Research Programs through the Ovarian Cancer Research Program (HT94252410269) (DP). This work was supported by the Dana-Farber/Harvard Cancer Center (DF/HCC) SPORE in Ovarian Cancer (DP). This work was also supported by the Massachusetts Life Science Council (MLSC) First Look Award (DP). The illustrations were created with BioRender.com.

## Author contribution

Conceptualization: M.C. and D.P.; methodology: M.C., E.T., C.C., HH.L., M.-C.M., P.M., L.G.; investigation, M.C., D.P., E.T., M.-C.M.; writing—original draft: M.C and D.P..; writing—review & editing: N/A; funding acquisition: D.P.; resources: D.P., M.C, HA.M, V.L., L.G., N.B; supervision, M.C. and D.P.

## Notes

### Competing Interest Statement

The authors have declared no competing interest.

## Bibliography

1. J. Huang, et al., Worldwide Burden, Risk Factors, and Temporal Trends of Ovarian Cancer: A Global Study. Cancers 14, 2230 (2022).

2. F. Bray, et al., Global cancer statistics 2022: GLOBOCAN estimates of incidence and mortality worldwide for 36 cancers in 185 countries. CA: A Cancer Journal for Clinicians 74, 229–263 (2024).

3. R. L. Siegel, K. D. Miller, N. S. Wagle, A. Jemal, Cancer statistics, 2023. CA Cancer J Clin 73, 17–48 (2023).

4. G. C. Jayson, E. C. Kohn, H. C. Kitchener, J. A. Ledermann, Ovarian cancer. The Lancet 384, 1376– 1388 (2014).

5. S. Pignata, S. C Cecere, A. Du Bois, P. Harter, F. Heitz, Treatment of recurrent ovarian cancer. Ann Oncol 28, viii51–viii56 (2017).

6. L. A. Baldwin, et al., Ten-Year Relative Survival for Epithelial Ovarian Cancer. Obstetrics & Gynecology 120, 612 (2012).

7. Cancer Genome Atlas Research Network, Integrated genomic analyses of ovarian carcinoma. Nature 474, 609–615 (2011).

8. K. P. Pennington, et al., Germline and somatic mutations in homologous recombination genes predict platinum response and survival in ovarian, fallopian tube, and peritoneal carcinomas. Clin Cancer Res 20, 764–775 (2014).

9. I. Vergote, et al., Clinical research in ovarian cancer: consensus recommendations from the Gynecologic Cancer InterGroup. Lancet Oncol 23, e374–e384 (2022).

10. J. Hamanishi, et al., Safety and Antitumor Activity of Anti-PD-1 Antibody, Nivolumab, in Patients With Platinum-Resistant Ovarian Cancer. J Clin Oncol 33, 4015–4022 (2015).

11. U. A. Matulonis, et al., Antitumor activity and safety of pembrolizumab in patients with advanced recurrent ovarian cancer: results from the phase II KEYNOTE-100 study. Annals of Oncology 30, 1080–1087 (2019).

12. M. L. Disis, et al., Efficacy and Safety of Avelumab for Patients With Recurrent or Refractory Ovarian Cancer. JAMA Oncol 5, 393–401 (2019).

13. B. Arneth, Tumor Microenvironment. *Medicina (Kaunas)* **56**, 15 (2019).

14. Y. Xiao, D. Yu, Tumor microenvironment as a therapeutic target in cancer. Pharmacol Ther 221, 107753 (2021).

15. M. Binnewies, et al., Understanding the tumor immune microenvironment (TIME) for effective therapy. Nat Med 24, 541–550 (2018).

16. S. Iyer, et al., Genetically Defined Syngeneic Mouse Models of Ovarian Cancer as Tools for the Discovery of Combination Immunotherapy. Cancer Discov 11, 384–407 (2021).

17. M. P. Hedger, W. R. Winnall, D. J. Phillips, D. M. de Kretser, The regulation and functions of activin and follistatin in inflammation and immunity. Vitam Horm 85, 255–297 (2011).

18. I. Morianos, G. Papadopoulou, M. Semitekolou, G. Xanthou, Activin-A in the regulation of immunity in health and disease. Journal of Autoimmunity 104, 102314 (2019).

19. P. Ren, et al., High serum levels of follistatin in patients with ovarian cancer. J Int Med Res 40, 877– 886 (2012).

20. A. Schneyer, et al., Differential actions of follistatin and follistatin-like 3. Molecular and Cellular Endocrinology 225, 25–28 (2004).

21. H. Li, et al., FSTL3 promotes tumor immune evasion and attenuates response to anti-PD1 therapy by stabilizing c-Myc in colorectal cancer. Cell Death Dis 15, 1–15 (2024).

22. Y. Sakamoto, et al., Determination of free follistatin levels in sera of normal subjects and patients with various diseases. Eur J Endocrinol 135, 345–351 (1996).

23. P. Florio, et al., High serum follistatin levels in women with ovarian endometriosis. Fertility and Sterility 88, S205 (2007).

24. X. Li, et al., FSTL3 is highly expressed in adipose tissue of individuals with overweight or obesity and is associated with inflammation. Obesity (Silver Spring) 31, 171–183 (2023).

25. S. Iyer, et al., Genetically defined syngeneic mouse models of ovarian cancer as tools for the discovery of combination immunotherapy. Cancer Discov (2020). 10.1158/2159-8290.CD-20-0818.

26. S. Muttukrishna, D. Tannetta, N. Groome, I. Sargent, Activin and follistatin in female reproduction. Mol Cell Endocrinol 225, 45–56 (2004).

27. T. Suzuki, et al., Role of epithelial-mesenchymal transition factor SNAI1 and its targets in ovarian cancer aggressiveness. jcmt 9, N/A-N/A (2023).

28. Y. Brown, S. Hua, P. S. Tanwar, Extracellular matrix in high-grade serous ovarian cancer: Advances in understanding of carcinogenesis and cancer biology. Matrix Biology 118, 16–46 (2023).

29. M. Zhang, Z. Chen, Y. Wang, H. Zhao, Y. Du, The Role of Cancer-Associated Fibroblasts in Ovarian Cancer. Cancers 14, 2637 (2022).

30. H. Fujimoto, et al., Tumor-associated fibrosis: a unique mechanism promoting ovarian cancer metastasis and peritoneal dissemination. Cancer Metastasis Rev 43, 1037–1053 (2024).

31. Y. Chhabra, A. T. Weeraratna, Fibroblasts in cancer: Unity in heterogeneity. Cell 186, 1580–1609 (2023).

32. X. Mao, et al., Crosstalk between cancer-associated fibroblasts and immune cells in the tumor microenvironment: new findings and future perspectives. Molecular Cancer 20, 131 (2021).

33. Y. Gao, et al., Cross-tissue human fibroblast atlas reveals myofibroblast subtypes with distinct roles in immune modulation. Cancer Cell 42, 1764–1783.e10 (2024).

34. S. A. Antar, N. A. Ashour, M. E. Marawan, A. A. Al-Karmalawy, Fibrosis: Types, Effects, Markers, Mechanisms for Disease Progression, and Its Relation with Oxidative Stress, Immunity, and Inflammation. International Journal of Molecular Sciences 24, 4004 (2023).

35. R. Lattouf, et al., Picrosirius red staining: a useful tool to appraise collagen networks in normal and pathological tissues. J Histochem Cytochem 62, 751–758 (2014).

36. The Cancer Genome Atlas Program (TCGA) - NCI. (2022). Available at: https://www.cancer.gov/ccg/research/genome-sequencing/tcga [Accessed 15 June 2023].

37. C. King, et al., LY2606368 Causes Replication Catastrophe and Antitumor Effects through CHK1- Dependent Mechanisms. Mol Cancer Ther 14, 2004–2013 (2015).

38. Prexasertib, a checkpoint kinase inhibitor: from preclinical data to clinical development - PubMed. Available at: https://pubmed.ncbi.nlm.nih.gov/31512029/ [Accessed 24 September 2024].

39. E. Giudice, et al., The CHK1 inhibitor prexasertib in BRCA wild-type platinum-resistant recurrent high-grade serous ovarian carcinoma: a phase 2 trial. Nat Commun 15, 2805 (2024).

40. A. Pawłowska, et al., Current Understanding on Why Ovarian Cancer Is Resistant to Immune Checkpoint Inhibitors. Int J Mol Sci 24, 10859 (2023).

41. P. Pandey, et al., New insights about the PDGF/PDGFR signaling pathway as a promising target to develop cancer therapeutic strategies. Biomedicine & Pharmacotherapy 161, 114491 (2023).

42. D. Peng, M. Fu, M. Wang, Y. Wei, X. Wei, Targeting TGF-β signal transduction for fibrosis and cancer therapy. Molecular Cancer 21, 104 (2022).

43. A. Ostman, PDGF receptors-mediators of autocrine tumor growth and regulators of tumor vasculature and stroma. Cytokine Growth Factor Rev 15, 275–286 (2004).

44. M. Pickup, S. Novitskiy, H. L. Moses, The roles of TGFβ in the tumour microenvironment. Nat Rev Cancer 13, 788–799 (2013).

45. L. Gorelik, R. A. Flavell, Transforming growth factor-beta in T-cell biology. Nat Rev Immunol 2, 46–53 (2002).

46. E. Batlle, J. Massagué, Transforming Growth Factor-β Signaling in Immunity and Cancer. Immunity 50, 924–940 (2019).

47. M. Chauvin, et al., Cancer-associated mesothelial cells are regulated by the anti-Müllerian hormone axis. Cell Reports 42, 112730 (2023).

48. H. Huang, et al., Mesothelial cell-derived antigen-presenting cancer-associated fibroblasts induce expansion of regulatory T cells in pancreatic cancer. Cancer Cell 40, 656–673.e7 (2022).

49. J. Qian, et al., Cancer-associated mesothelial cells promote ovarian cancer chemoresistance through paracrine osteopontin signaling. J Clin Invest 131, 146186 (2021).

50. A. Cho, V. M. Howell, E. K. Colvin, The Extracellular Matrix in Epithelial Ovarian Cancer – A Piece of a Puzzle. Frontiers in Oncology 5, 245 (2015).

51. C. Chandler, T. Liu, R. Buckanovich, L. G. Coffman, The double edge sword of fibrosis in cancer. Translational Research 209, 55–67 (2019).

52. H. Jiang, S. Hegde, D. G. DeNardo, Tumor-associated fibrosis as a regulator of tumor immunity and response to immunotherapy. Cancer Immunology, Immunotherapy : CII 66, 1037 (2017).

53. X. Meng, et al., FSTL3 is associated with prognosis and immune cell infiltration in lung adenocarcinoma. J Cancer Res Clin Oncol 150, 17 (2024).

54. C.-H. Li, et al., The cytoplasmic expression of FSTL3 correlates with colorectal cancer progression, metastasis status and prognosis. J Cell Mol Med 27, 672–686 (2023).

55. H. Zhu, et al., Elevation of ADAM12 facilitates tumor progression by enhancing metastasis and immune infiltration in gastric cancer. Int J Oncol 60, 51 (2022).

56. Y. Liu, et al., Bioinformatic Analyses and Experimental Verification Reveal that High FSTL3 Expression Promotes EMT via Fibronectin-1/α5β1 Interaction in Colorectal Cancer. Front Mol Biosci 8, 762924 (2021).

57. Y.-J. Liu, et al., FSTL3 is a Prognostic Biomarker in Gastric Cancer and is Correlated with M2 Macrophage Infiltration. Onco Targets Ther 14, 4099–4117 (2021).

58. C. Yang, F. Cao, S. Huang, Y. Zheng, Follistatin-Like 3 Correlates With Lymph Node Metastasis and Serves as a Biomarker of Extracellular Matrix Remodeling in Colorectal Cancer. Front Immunol 12, 717505 (2021).

59. The Human Protein Atlas. Available at: https://www.proteinatlas.org/ [Accessed 4 December 2024].

60. M. Uhlen, et al., A pathology atlas of the human cancer transcriptome. Science 357, eaan2507 (2017).

61. UCSC Xena. Available at: https://xenabrowser.net/datapages/?dataset=TCGA.OV.sampleMap%2FHiSeqV2_exon&host= https://tcga.xenahubs.net&removeHub= https://xena.treehouse.gi.ucsc.edu%3A443 [Accessed 20 November 2024].

